# Telomerase and Alternative Lengthening of Telomeres coexist in the regenerating zebrafish caudal fins

**DOI:** 10.1101/2021.11.15.468592

**Authors:** Elena Martínez-Balsalobre, Monique Anchelin-Flageul, Francisca Alcaraz-Pérez, Jesús García-Castillo, David Hernández-Silva, Maria C Mione, Victoriano Mulero, María L. Cayuela

## Abstract

Telomeres are essential for chromosome protection and genomic stability, and telomerase function is critical to organ homeostasis. Zebrafish has become a useful vertebrate model for understanding the cellular and molecular mechanisms of regeneration. The regeneration capacity of the caudal fin of wild-type zebrafish is not affected by repetitive amputation, but the behavior of telomeres during this process has not yet been studied. In this study, the regeneration process was characterized in a telomerase deficient zebrafish model. Moreover, the regenerative capacity after repetitive amputations and at different ages was studied. Regenerative efficiency decreases with aging in all genotypes and surprisingly, telomere length is maintained even in telomerase deficient genotypes. Our results suggest that telomere length can be maintained by the regenerating cells through the recombination-mediated Alternative Lengthening of Telomeres (ALT) pathway, which is likely to support high rates of cell proliferation during the tailfin regeneration process. As far as we know, this is the first animal model to study ALT mechanism in regeneration, which opens a wealth of possibilities to study new treatments of ALT dependent processes.

## INTRODUCTION

Tissue regeneration is an evolutionary conserved response to injury (Morrison et al, 2006) and while all animals regenerate some of their tissues by physiological turnover, only few can regenerate appendages. Zebrafish (*Danio rerio*) is one of such organisms, able to regenerate retina, fins, heart, spinal cord and other tissues even in advanced ages (Becker et al, 1997; Poss et al, 2003; Poss et al, 2002; Reimschuessel, 2001; Rowlerson et al, 1997). Because of its regenerative ability, its simple but relevant anatomy, *in vivo* imaging capability and genetic advantages, zebrafish have become a useful vertebrate model for understanding the cellular and molecular mechanisms of regeneration (Goldsmith & Jobin, 2012). Caudal fin is the most convenient tissue for regenerative studies due to its easy handling and fast regeneration. Adult zebrafish regenerate their caudal fin within fourteen days after amputation (Anchelin et al, 2011; Poss et al, 2003).

Several groups have investigated cells and genetic signalling pathways regulating blastema formation (Wehner & Weidinger, 2015). In addition, the regeneration limit of the zebrafish caudal fin was recently investigated (Azevedo et al, 2011; Shao et al, 2011), however, there are very few studies about the implication of telomeres and telomerase in this process. Moreover, the zebrafish has constitutively abundant telomerase activity in somatic tissues from embryos to aged adults (Anchelin et al, 2011; McChesney et al, 2005). Notably, a study on various tissues from aquatic species including the zebrafish suggests that telomerase may be important for tissue renewal and regeneration after injury rather than for overall organism longevity (Elmore et al, 2008). Our previous studies about behaviour of telomeres and telomerase during the regeneration process revealed a direct relationship between telomerase expression, telomere length and efficiency of tissue regeneration in wild-type zebrafish caudal fin (Anchelin et al, 2011). In addition, characterization of the telomerase-deficient zebrafish suggest that telomerase function is crucial for organ homeostasis in zebrafish (Anchelin et al, 2013; Henriques et al, 2013), as occurs in mouse (Blasco et al, 1997).

The aim of this study was to clarify the role of telomerase during the caudal fin regenerating process. We confirmed the outstanding and almost unlimited caudal fin regeneration capability of zebrafish, which only decrease in aged animals, also in the case of telomerase-deficient genotype. Moreover, we found the telomere length is maintained even in telomerase-deficient genotypes suggesting the involvement of Alternative Lengthening of Telomeres (ALT) pathway.

## RESULTS

### Caudal fin regeneration is affected by aging

It is known that after a single excision of the zebrafish caudal fin, a regenerative process is activated and it takes approximately two weeks to fully regenerate all the tissues and structures that compose a functional fin (Akimenko et al, 2003; Poss et al, 2003). Several studies show that consecutive repeated amputations of zebrafish caudal fin do not reduce its regeneration capacity (Azevedo et al, 2011; Shao et al, 2011); however, changes in telomere length during the regenerative process have not been studied.

To further investigate the role of telomerase during this regenerating process, *tert* ^*+/+*,^ *tert* ^*+/−*^ and *tert* ^*−/−*^ specimens from the same genetic background, maintained in the same laboratory conditions, were used for regeneration assays after a single amputation or after repeated amputations.

Zebrafish of all three genotypes and at different life stages (4, 8 and 11 months) were amputated and their regeneration was monitored (Fig. 1A). All genotypes can regenerate the caudal fin. The 8-month fish regeneration curve (Fig. 1B) showed that *tert* ^*+/+*^ regenerates its entire tail fin at 16 days post-amputation (dpa), faster than *tert* ^*+/-*^ and *tert* ^*-/-*^ zebrafish (at 19 and 22 dpa, respectively). Furthermore, *tert* ^*+/-*^ and *tert* ^*-/-*^ zebrafish need more time to reach 50% of their caudal fin regeneration than wild-type zebrafish at all ages (Fig. 1C), and this difference is significant at 11 months of age, when fish without telomerase are considered to be aged (Anchelin et al, 2013).

**Figure 1.**
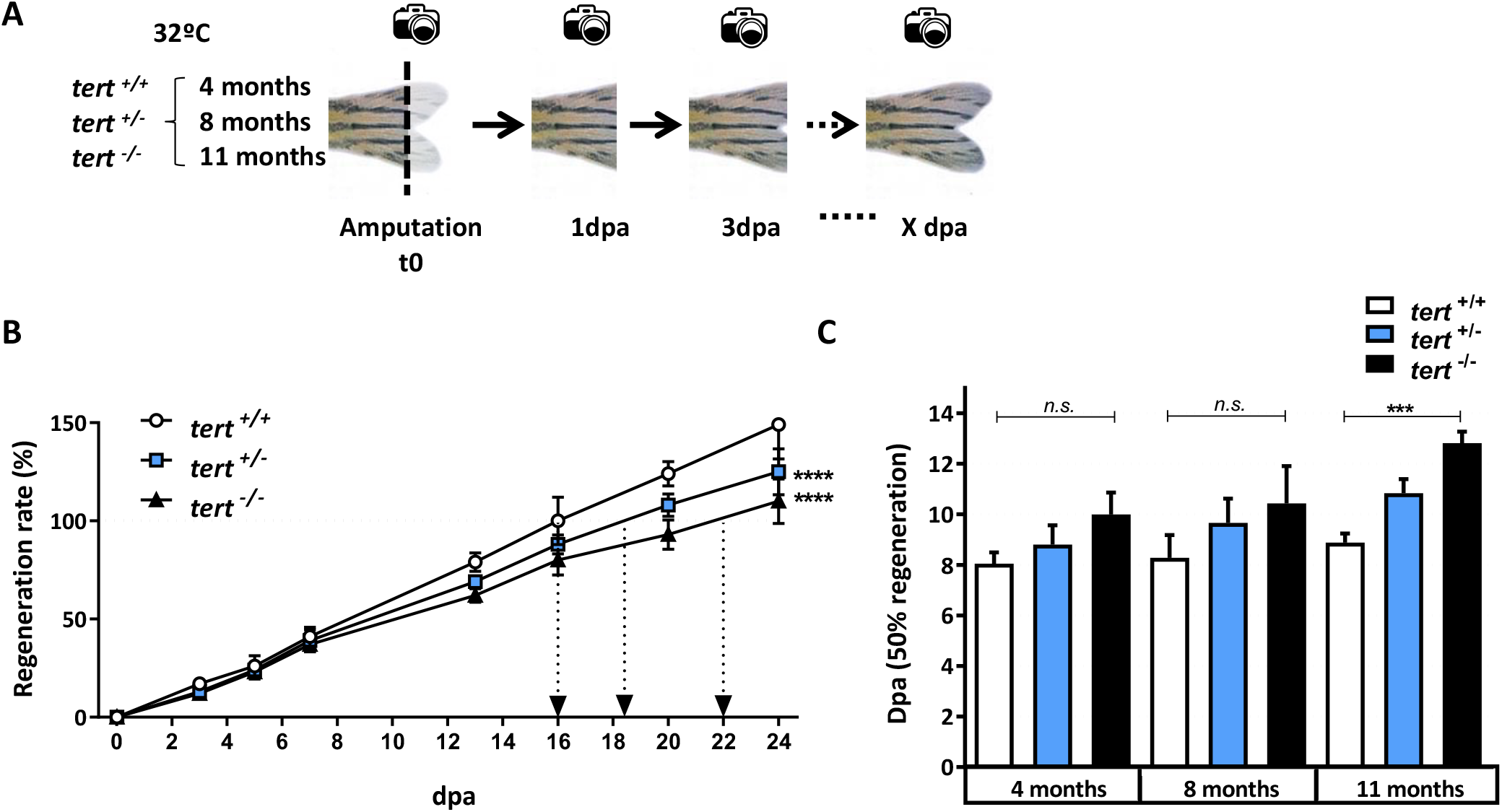
Telomerase genotypes regenerate with different rate. **A**, Experimental design of a regeneration assay with a single amputation and growth monitoring, at three different stages of life (4, 8, and 11 month-old) in *tert* ^*+/+*^, *tert* ^*+/-*^and *tert* ^*-/-*^zebrafish. **B**, Caudal fin regeneration curve of 8 month-old zebrafish from the three genotypes. The arrows indicate 100% regeneration of the tail fin. n=6. **C**, Days to reach 50% of the caudal fin regeneration at three different ages and from the three telomerase genotypes. n=6. All data are mean + s.e.m. ***P< 0.001 and **** P< 0.0001 for two-way analysis of variance (ANOVA) plus Dunnett’s post-test.

Since the potencial for caudal fin regeneration has been shown to be neither reduced by consecutive clips nor their telomere length reduced by three consecutive clips (Anchelin et al, 2011; Azevedo et al, 2011; Shao et al, 2011) we explored the regeneration process in telomerase deficient zebrafish. We performed a consecutive repeated amputation experiment using “young adult fish” (4 month-old) and “old adult fish” (11 month-old) from the three genotypes (Fig. 2A). The resulted regeneration curve showed that after 12th repeated consecutive amputations, with 7 days intervals, the young zebrafish from the three different genotypes are able to regenerate their caudal fin at a similar regeneration rate (Fig. 2B). However, in the case of the old adult fish, *tert* ^+/−^ and *tert* ^−/−^ zebrafish showed a lower, statistically significant reduction of regeneration rate compared with their wild-type *tert* ^+/+^ (Fig. 2C) counterpart, which continues to show a high regeneration rate, although slower than their younger siblings.

**Figure 2.**
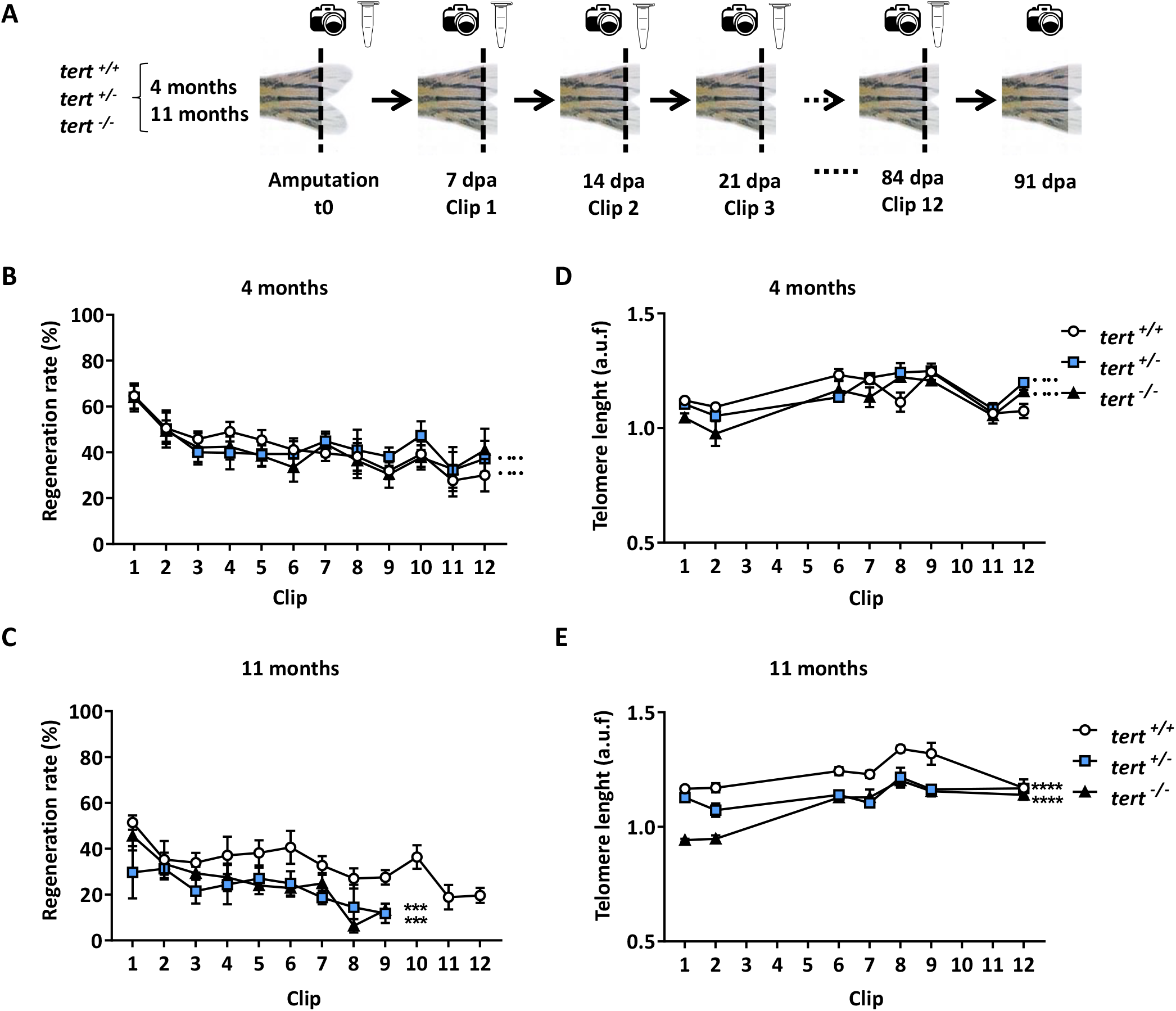
Telomere length is maintained during consecutive amputations in telomerase deficient zebrafish. **A**, Experimental design of a regeneration assay with 12 consecutive amputations every 7 days and growth monitoring, in *tert* ^*+/+*^, *tert* ^*+/-*^and *tert* ^*-/-*^genotypes, at 4 and 11 months of age. **B**, Curves of the regeneration rates after 12 consecutive clips in the three telomerase genotypes at 4 months age. n=6 **C**, Curves of the regeneration rates after 12 consecutive clips in the three telomerase genotypes at 11 months age. n=6 **D**, Measurement of telomere length by Flow-FISH of regenerative fin tissue cells in all three telomerase genotypes at 4 months of age. **E**, Measurement of telomere length by Flow-FISH of regenerative fin tissue cells in all three telomerase genotypes at 11 months of age. Data are average of at least 2 independent experiments. All data are mean + s.e.m. ***P< 0.001 and **** P< 0.0001 for two-way analysis of variance (ANOVA) plus Dunnett’s post-test.

### Telomere length is maintained during consecutive amputations assay in all three telomerase genotypes

It has already been published and confirmed here, that the regenerative capacity of zebrafish tail fin is unlimited. Surprisingly, this regenerative capacity is not diminished in telomerase deficient genotypes (*tert* ^+/-^ and *tert* ^-/-^), therefore we decided to assess the dynamic of telomere length. Fin regenerative tissue was collected from each group at different clips along the regeneration assay and the telomere length was measured by Flow-FISH. Telomere length was maintained in the young zebrafish group even in telomerase deficient genotypes (Fig. 2D). However, in old zebrafish the telomere lenght decreased in *tert* ^*+/-*^ and *tert* ^*-/-*^ (Fig. 2E), but not dramatically as expected.

These results suggest that telomerase is not essential for the regeneration of the zebrafish caudal fin and lead us to further investigate the mechanism by which these cells are able to maintain their telomere length throughout a repeated amputations assay, which means a continual renewal of tissues with high cell proliferation rates.

### ALT is involved in telomere length maintenance during the regeneration process

Telomerase is the main telomere maintenance mechanism, but in about 10% of tumors cells, telomere length is maintained by the Alternative Lengthening of Telomeres (ALT) mechanism (Conomos et al, 2013; Draskovic & Londono Vallejo, 2013). This mechanism is based on homologous recombination and homology-directed telomere synthesis (Pickett & Reddel, 2015). ALT cells show a number of unusual characteristics; one of the most striking is an abundance of DNA with telomeric sequences separate from the chromosomes. This extrachromosomal telomeric DNA takes many forms, including predominantly double-stranded telomeric circles (referred to as C-circles, as it mostly involves the C-rich strand) (Henson et al, 2009).

Therefore, we set out to detect and quantify circular telomeric DNA in the cells obtained from the caudal fin regenerative portion using the quantitative and sensitive CCassay (Lau et al, 2013) (Fig. 3A). At time zero there was no significant differences between wild-type and telomerase deficient fish in C-circles amounts. However, we observed a statistically significant increase above 2 (a value considered indicative of C-circles abundance,(Henson et al, 2009)) of C-circles at 48 hours post amputation (hpa) in the cells from the telomerase-deficient zebrafish compared with no changes observed in the cells of the wild-type zebrafish. As controls for C-circle levels, a telomerase cell line (HeLa) and an ALT cell line (SAOS-2) were used (Fig. S1A)

**Figure 3.**
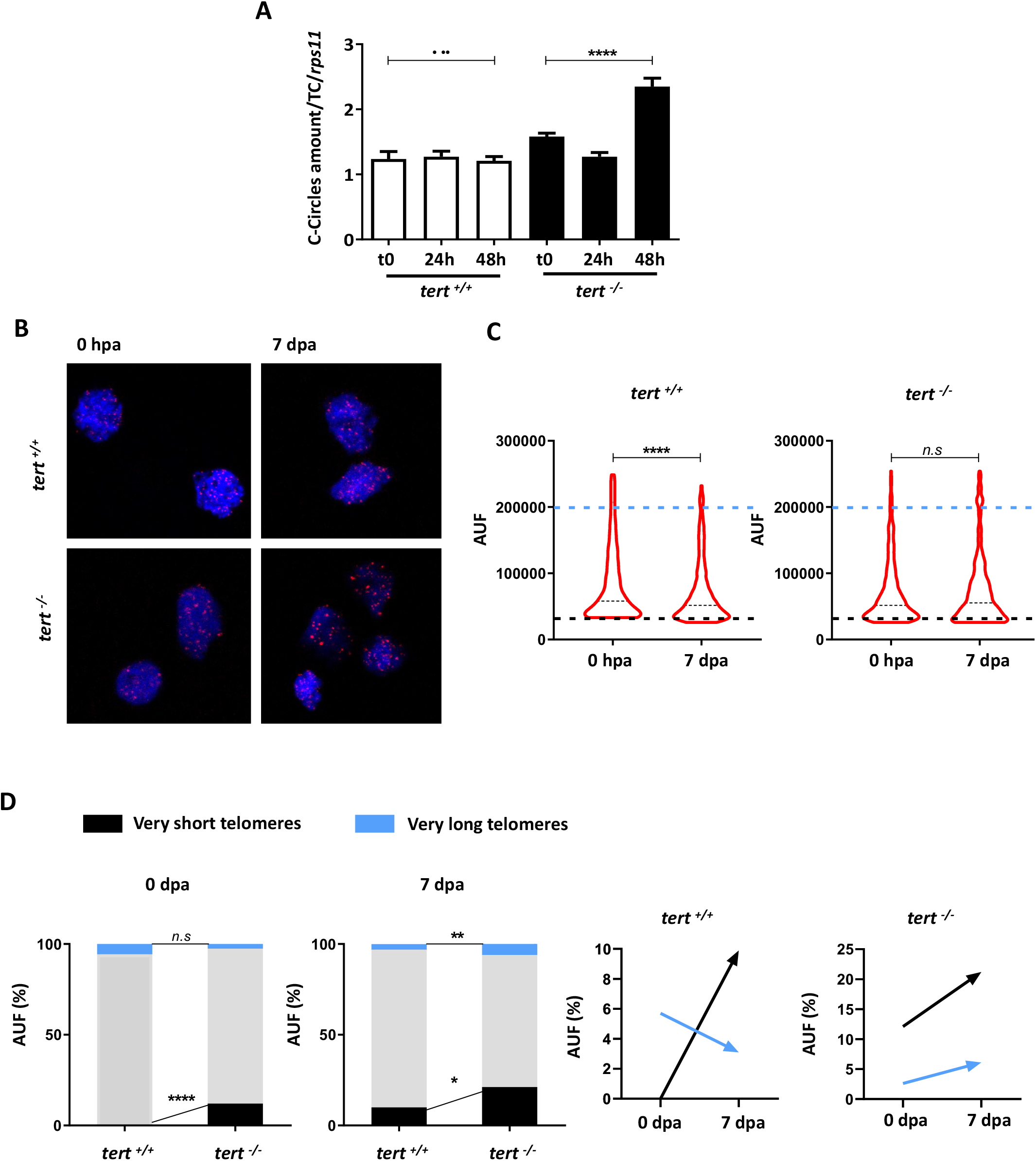
Telomerase deficient zebrafish exhibit C-circles and heterogeneous telomere length in regenerative tissue. **A**, C-circles abundance in regenerative tissue from wildtype and telomerase deficient adult zebrafish at 24 and 48 hpa. **B**, Representative images of Q-FISH in blastema cells from *tert* ^*+/+*^ and *tert* ^*-/-*^genotypes at time 0 hpa and 7 dpa. **C**, Telomeric fluorescence values (arbitrary units of fluorescence, AUF) of regenerative cells from *tert* ^*+/+*^ and *tert* ^*-/-*^adult zebrafish at two times. **D**, Telomeric fluorescence frequency of blastema cells from *tert* ^*+/+*^ and *tert* ^*-/-*^adult zebrafish at 0 dpa and 7 dpa. Very short telomeres have a higher fluorescence of 200,000 AUF and very short telomeres have a lower fluorescence of 30,000 AUF. Data are average of at least 2 independent experiments. All data are mean + s.e.m. *n*.*s*. not significant. **** P< 0.0001 for, One-way analysis of variance (ANOVA) plus Dunnett’s post-test (A) and t-test Student (C, D).

Other characteristics of ALT cells include highly heterogeneous telomere lengths (ranging from undetectable to extremely long), and rapid changes in telomere length (Cesare & Griffith, 2004). To visualize zebrafish telomeres during regeneration, a quantitative FISH (q-FISH) was performed to calculate relative length of telomere within the blastema interphase cells before amputation and 7 days post amputations (Fig. 3B-D). We observed a decrease in telomere length after the proliferation process leading to regeneration starts in wild-type zebrafish but not in fish of *tert* ^*-/-*^ line, where we did not detected changes in the average telomeric length (Fig. 3C). However, we observed a higher (Fig. 3D) telomeric heterogeneity in *tert* ^*-/-*^ than in *tert* ^*+/+*^. A higher percentage of short (21.1%) and long telomeres (6.1%) in telomerase deficient zebrafish 7 days post amputation was detected compared with percentage of short (3.1%) and long telomere (9.9%) before amputation (Fig 3D).

Since ALT is based on homologous recombination between telomeres, a number of recombination proteins have been shown by genetic analyses to be necessary for telomere maintenance in ALT cells (Martinez et al, 2017), as the MRN complex (MRE11a; RAD 50 and NBS1) (Zhong et al, 2007) and other DNA binding proteins, including the serine/threonine protein kinase, ATR (Collis et al, 2008). On the other hand, chromatin in ALT cells seems to be more relaxed than the one of telomerase–positive cells. Interestengly, mutations in the genes encoding ATRX (α-thalassemia/mental retardation syndrome X-linked) and DAXX (death domain-associated protein), members of chromatin remodeling complex active at telomeres, are strongly associated with ALT and may be involved in ALT suppression, while their loss is associated with ALT activation (Amorim et al, 2016; Ren et al, 2018; Yost et al, 2019). Therefore, we decided to analyze the mRNA levels of several genes, which may be required to activate or prevent ALT during the regeneration process. These genes were quantified in the regenerated tissue at 24 and 48 hpa. *nbs1* and *atr* mRNA levels increased in telomerase deficient fish at 48 hpa compared to time 0 (t0) (Fig. 4A, 4B), whereas *atrx* and *daxx* expression decreased (Fig. 4C, 4D) at 24 and 48 hpa, in agreement with published data on ALT in cells (Amorim et al, 2016; Ren et al, 2018; Yost et al, 2019).

**Figure 4.**
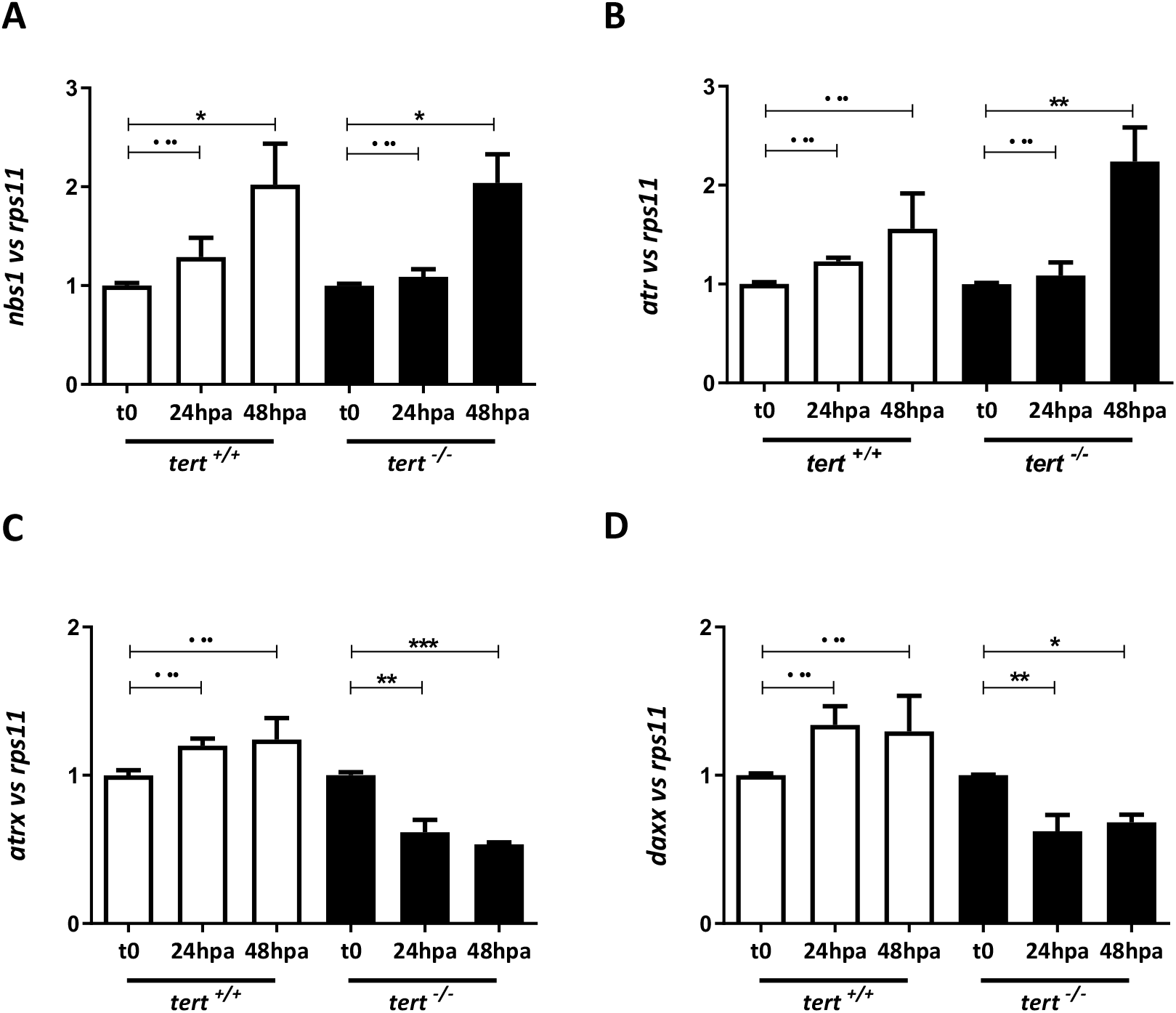
ALT mechanism is activated in regenerative tissue of telomerase deficient zebrafish. **A**, *nbs1* expression normalized to *rps11* expression in regenerative tissue from wildtype and telomerase deficient adult zebrafish at 24 and 48 hpa. **B**, *atr* expression normalized to *rps11* in blastema tissue from *tert* ^*+/+*^ and *tert* ^*-/-*^adult zebrafish at 24 and 48 hpa. **C**, *atrx* expression normalized to *rps11* in regenerative tissue from *tert* ^*+/+*^ and *tert* ^*-/-*^adult zebrafish at 24 and 48 hpa. **D**, *daxx* expression normalized to *rps11* in regenerative tissue from *tert* ^*+/+*^ and *tert* ^*-/-*^adult zebrafish at 24 and 48 hpa. Data are average of at least 2 independent experiments. All data are mean + s.e.m. *n*.*s*. not significant. *P< 0.05, **P< 0.01 and *** P< 0.001 for One-way analysis of variance (ANOVA) plus Dunnett’s post-test.

Together, these data demonstrate the involvement of ALT in the regenerative process in telomerase deficient fish.

### Inhibition of *nbs1* and *atr* genes affects caudal fin regeneration

NBS1 protein belonging to MNR recombination protein complex (MRE11–RAD50–NBS1) implicated in homologue recombination process is required for both ALT phenotype development and extrachromosomal telomeric circles formation. Its inhibition decreased numbers of ALT-associated promyelocytic leukemia bodies and decreased telomere length (Compton et al, 2007; Zhong et al, 2007). In the same way, ATR inhibition, which kinase activity is activated by NBS, disrupt ALT and selectively kill ALT cells *in vitro* (Flynn et al, 2015).

To further investigate whether telomere length was maintained through ALT mechanism during the regeneration process, *nbs1* and *atr* were downregulated by injection of a vivo fluorescent morfolino, using a standard morpholino as negative control. Vivo morfolinos are oligomers used to knockdown a gene as the commonly used ATG or splice-blocking morpholinos, but modified for *in vivo* experiments. The injection was performed in the dorsal area of 48 hpa regenerated tissue of the caudal fin, followed by electroporation of the injected zone to facilitate the morpholino entry into the cells. 48 hours post-injection (hpi) the regeneration rate was calculated, using the ventral area as an internal control (Fig. 5A).

**Figure 5.**
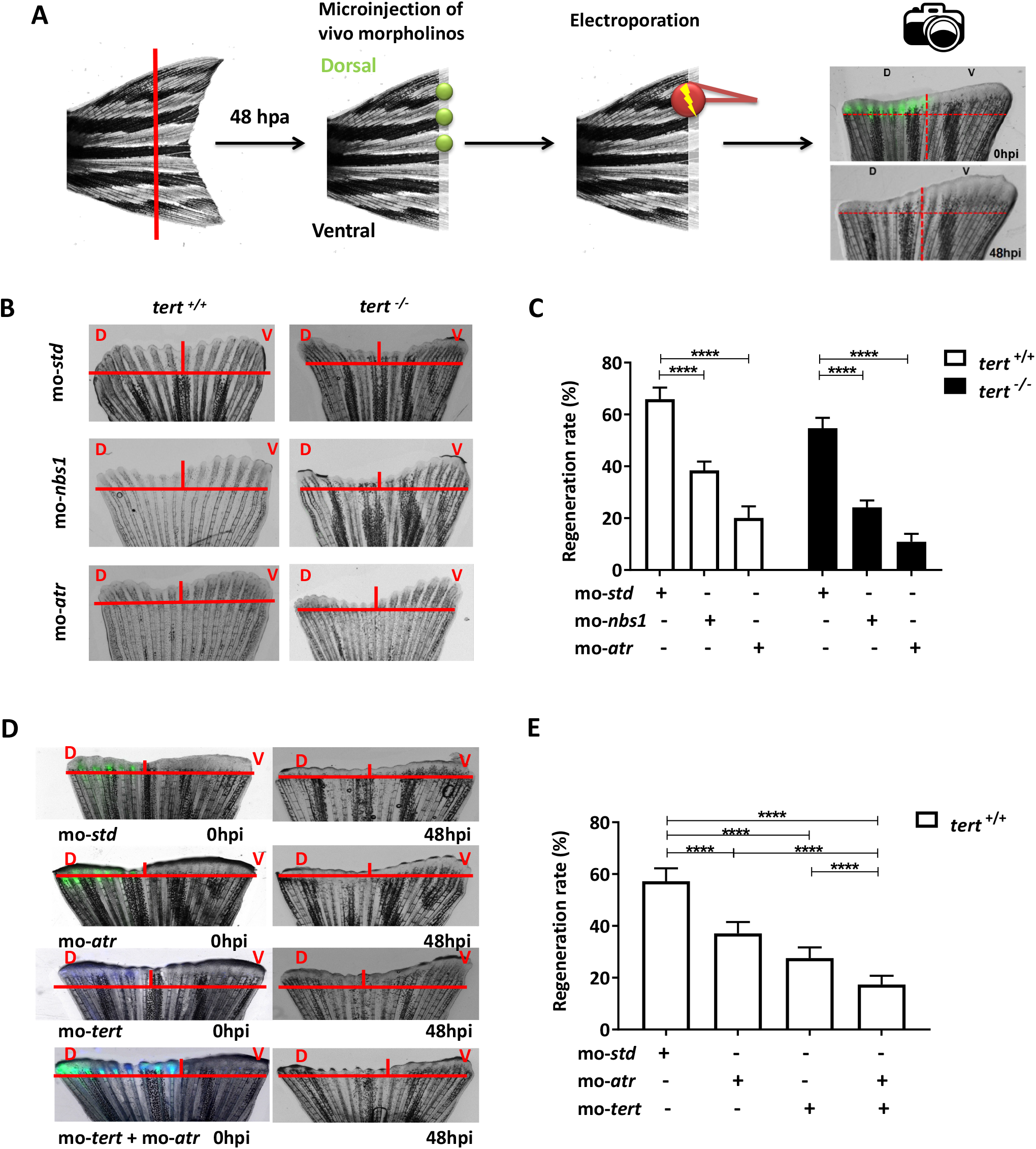
Inhibition of *nbs1* and *atr* affects adult caudal fin regeneration. **A**, Schematic representation of vivo fluorescent morpholino microinjection into dorsal area of caudal fin after 48 hpa. Microinjected area is then electroporated. Ventral area is used as a control of normal regeneration rate. After 48 hpi, the regeneration rate was calculated. **B**, Representative pictures of caudal fin regeneration after 48h post-morpholino microinjection. **C**, Regeneration rate of caudal fin in *tert* ^*+/+*^ and *tert* ^*-/-*^genotypes with *nbs1* and *atr* knockdown. n=7 **D**, Representative images of caudal fin at 0 hpi of vivo morpholinos (standard and *atr* in green and *tert* in blue) in the dorsal half (D), and after 48 hpi. (V) uninjected ventral half. **E**, Regeneration rate of caudal fin in wildtype fish with *atr* and/or *tert* knockdown. n=10. All data are mean + s.e.m. **** P< 0.0001 for One-way analysis of variance (ANOVA) plus Sidak’s post-test.

Our results showed that inhibition of *nbs1* and *atr* genes blocks regeneration in both genotypes, more remarkably in *tert* ^*-/-*^ compared to *tert* ^*+/+*^ zebrafish (Fig. 5B and 5C).

To understand the contribution of ALT and Telomerase to the regenerative process, a regeneration experiment similar to the previous one was carried out in wild-type zebrafish. *atr* (to inhibit ALT) and *tert* (to inhibit telomerase) vivo fluorescent morfolinos were microinjected into the dorsal area of 48 hpa regenerated tissue (Fig. 5D). Regeneration was blocked also in these experimental conditions. The result (Fig. 5E) confirms that blocking both telomere maintaining mechanism (telomerase-dependent or ALT) decreased the regeneration capacity of zebrafish caudal fin. Interestently a synergistic effect was observed when both mechanisms are inhibited. Therefore, telomerase and ALT mechanisms are involved in zebrafish fin regeneration and when fish do not have one of these mechanisms, they can use the other to regenerate.

### Pharmacological inhibition of ATR in adults and larvae affects regeneration

As mentioned before, ALT is a recombination-based telomere maintenance mechanism utilized by 10-15% of human cancers (tumors of mesenchymal origin and often portends a poor prognosis). We decided to test pharmacological inhibition to target ALT in this model, since ALT is an attractive target for cancer therapy.

For this purpose, ATR Inhibitor IV (Calbiochem), an ATR-selective inhibitor, was used to study the effect of pharmacological inhibition of ALT (Flynn et al, 2015). Zebrasfish larvae treated with different concentration of ATR Inhibitor IV (1 µM to 50 µM) for 24 hours were assayed in a western blot against ATR/ATM substrates (#2851 Cell Signaling) to check the inhibitory effect of the treatment. The lower concentration did not affect the phosphorylation of ATR/ATM subtrates after 24h treatment in zebrafish larvae (data not show) however 10 µM and 50 µM showed similar inhibitory effects (Fig. S2A, S2B). Therefore the 10 µM concentration was used for our studies.

The effect of ATR inhibition during the fin regeneration process was analyzed by a regeneration assay on wild-type and *tert* deficient adult zebrafish (Fig. 6A). Regeneration rate was measure at 24 and 48 hours post amputation (hpa). Treatment with ATR inhibitor IV decreased the regenerative capacity of wild-type and *tert* deficient fish (Fig 6B and 6C).

**Figure 6.**
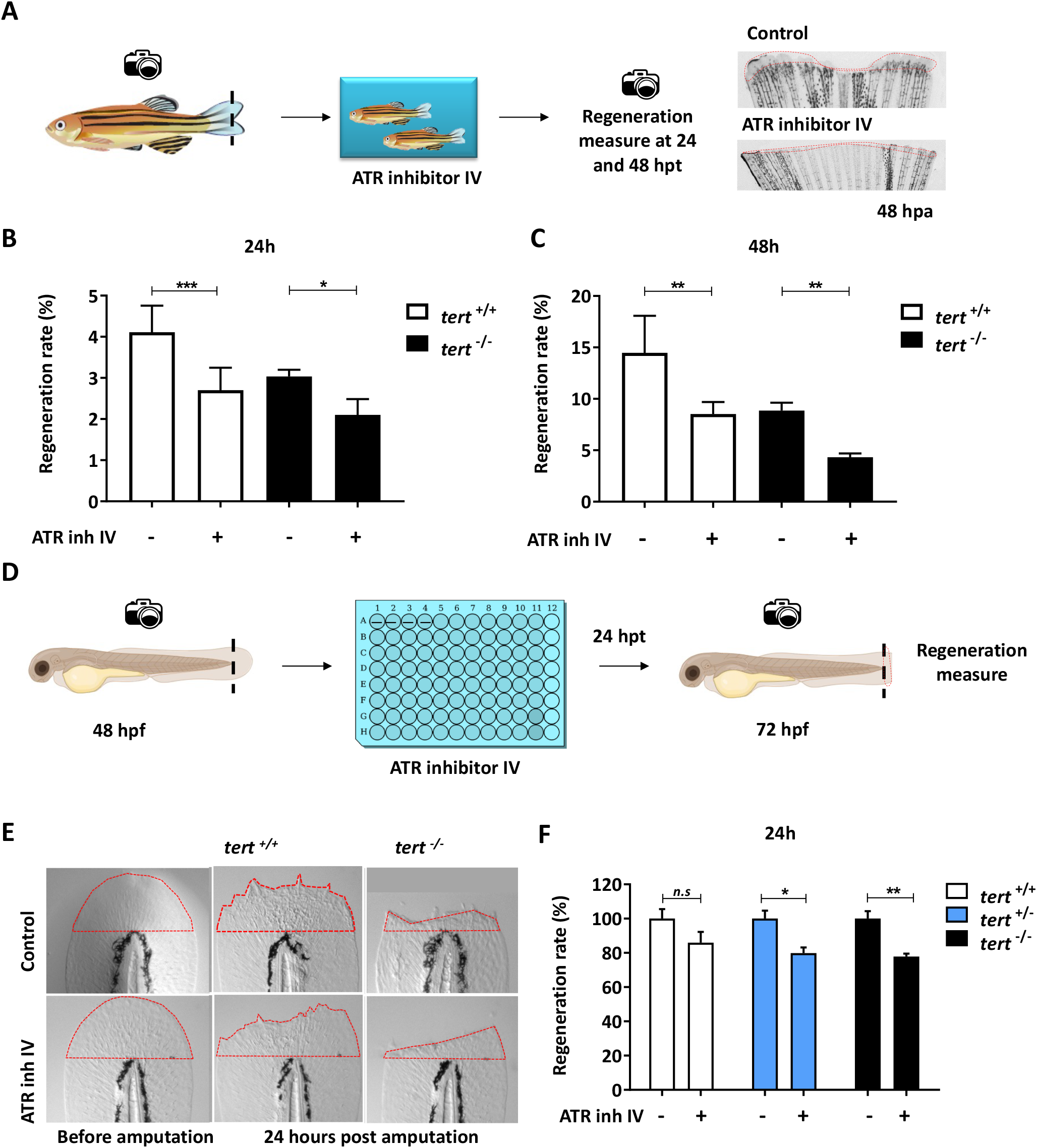
Pharmacological inhibition of ATR in larvae and adults affects regeneration. **A**, Experimental design of regeneration assay in adult zebrafish. ATR inhibitor IV is administered to adult fish tanks immediately after tail fin amputation and regenerative capacity is tested at 24 and 48 hpa, compared with untreated controls. **B**, Regeneration rate of caudal fin in adults at 24 hpa. n=6. **C**, Regeneration rate of caudal fin in adults at 48 hpa. n=6. **D**, Experimental design of regeneration assay in 48 hpf zebrafish larvae treated with ATR inhibitor IV after amputation for 24 hours and posterior tissue regeneration measure. **E**, Representative images of tail regeneration in *tert* ^*+/+*^ and *tert* ^*-/-*^larvae treated with ATR inhibitor IV and its respective controls. **F**, Regeneration rate of tail regeneration in *tert* ^*+/+*,^ *tert* ^*+/-*^and *tert* ^*-/-*^larvae treated with ATR inhibitor IV (ATRinh IV) and untreated controls. n≥100 larvae per group. All data are mean + s.e.m. *P< 0.05, **P< 0.01 and *** P< 0.001 for One-way analysis of variance (ANOVA) plus Sidak’s post-test.

Since zebrafish larvae are a powerfull tool for high-throuput screening we developed a regeneration assay on larvae for pharmacological screening. We treated 48 hpf zebrafish larvae with 10 µM ATR inhibitor IV by immersion immediately after tail amputation (Fig. 6D). 24 hours later (72 hpf larvae) the regeneration area was measured. As expected, the larvae treated with ATR inhibitor showed a smaller regenerated area compared to untreated larvae, although only statistically significant for telomerase-deficient genotypes (Fig. 6E, 6F).

Therefore, *atr* inhibition decreased the regenerative capacity of the zebrafish’s fin, more significantly in telomerase deficient fish. This means that the fin regeneration assay in *tert* ^-/-^ zebrafish could be used as a fast and simple model to identify drugs that inhibit ALT activity.

## DISCUSSION

Regeneration is a complex process of cell renewal, restoration and tissue growth. High proliferation rates are accompanied by telomerase activity to maintain telomere length to sustain replication and cell division.

Zebrafish has become a powerful model for investigating regenerative processes because of its capacity to completely repair several organs after injury.

An amputated fin ray is covered within the first several hours by epidermis, and within one to two days, a regeneration blastema forms. The blastema is a proliferative mass of morphologically similar cells, formed through disorganization and distal migration of fibroblasts and osteoblasts (or scleroblasts) proximal to the amputation plane (Gemberling et al, 2013). High proliferation rates accompainied by telomerase attrition required telomere maintenance to sustain replication and cell division. Howewer the involvement of telomerase and telomeric length behaviour have not been caracterized during caudal fin regeneration.

Our results confirmed the notable regeneration capacity of the zebrafish after a single or successive fin amputation (Anchelin et al, 2011; Azevedo et al, 2011; Shao et al, 2011), even in telomerase deficient zebrafish.

Surprisingly, telomerase-deficient zebrafish maintained its regenerative ability in both 4-month-old young fish and in prematurely aged 11-month fish. Although the regenerative capacity was not lost in old fish, it was reduced both after a single clip and by successive clips. Unexpectedly, they were able to keep the length of their telomeres constant after successive clips, suggesting that althought telomerase has an important role in regeneration, the participation of an alternative mechanism of telomere lengthening in zebrafish telomerase deficiency is essential.

In the case of telomerase deficiency, homologous recombination constitutes an alternative method (ALT ‘alternative lengthening of telomeres’) to maintain telomere DNA. Rolling-circle replication may use extrachromosomal t-circles, as has been shown in human ALT cells and in a wide variety of organisms including yeasts, higher plants and *Xenopus laevis*, to elongate telomere. From an evolutionary perspective, this widespread occurrence of t-circles may not only represent a back-up in the event of telomere dysfunction, but may be the primordial systems of telomere maintenance (Fajkus et al, 2005).

The presence of extrachromosomal telomeric circles, proposed to result from homologous recombination events at the telomeres, has been used as a marker of ALT cells. At 24 hpa, we detected telomeric DNA circles in the regenerative cells from the wild-type and the mutant zebrafish *tert* ^*-/-*^, although the amount of telomeric DNA circles is significantly higher in the mutant fish at 48 hpa, pointing out ALT has been activated in these cells.

Moreover, other charactesistics of ALT cells include a highly heterogeneous chromosomal telomere length (Murnane et al, 1994), which was observed after 7 dpa in *tert* ^*-/-*^, where the percentage of much shortened telomeres and very long telomeres were duplicated. Contrary to was observed in control fish, where the percentage of very short telomeres increased and the percentage of very long telomeres decreased.

Although the mechanism and causes of ALT are still not well-known, we wanted to study the expression of proteins involved in ALT during the regeneration process, including proteins involved in DNA repair as NBS1, also a critical regulator of recombination recruited by RPA as ATR complex and chromatin remodeling complex DAXX and/or ATRX wich mutations is associated with ALT (Hu et al, 2016; Lovejoy et al, 2012; Napier et al, 2015). Curiosly we observed an increased expresssion of ALT activator proteins and a decrease in ALT inhibitor proteins in telomerase deficient zebrafish, suggesting that the main players of ALT and their mechanisms are conserved during evolution.

Moreover, by inhibiting *nbs1* or *atr* by vivo morpholinos or ATR pharmacologically, thus preventing the maintenance of telomere by ALT, a lower regeneration efficiency was observed during caudal fin regeneration in wild-type and telomerase deficient zebrafish, although the effect is always more evident in *tert* ^*-/-*^ genotype. The regenerating cells in *tert* ^*-/-*^ present all the characteristics of ALT positive cells, such as telomeric heterogeneity, presence of extrachromosomal telomeric circles and over-expression of genes involved in homologous recombination.

Finally, *tert* and *atr* knockdown showed an impaired regeneration rate, which showed sinergy when both proteins were knocked down at the same time during the regeneration process. Together, our results lead us to conclude that in the absence of telomerase, telomere length is maintained through a homologous recombination mechanism during the early phases of regeneration. By contrast, in the wild-type zebrafish, the two mechanisms (telomerase-dependent and telomerase-independent) could coexist and participate to the maintenance of telomeres although ALT inhibition causes less severe effect than telomerase inhibition. During the zebrafish aging, dysfunctional telomeres could lead to the emergence of telomere recombination and so to ALT activation without hamper telomerase function (Brault & Autexier, 2011). The inhibition of both mechanisms practically prevents the regeneration of the tail fin (Fig. 7). Coexistence of these two mechanisms has been described in cancer *ex-vivo* and *in vitro* (De Vitis et al, 2018). The capability of some cancer cells *ex-vivo* to “switch” from one mechanism to the other is important, in a particular way for the application of dual targeted (anti-telomerase and anti-ALT) therapy to avoid drug resistance and will promote better therapies. Importantly zebrafish tail regeneration provides the first *in vivo* model to study both telomere maintenance mechanism and their coexistence.

**Figure 7.**
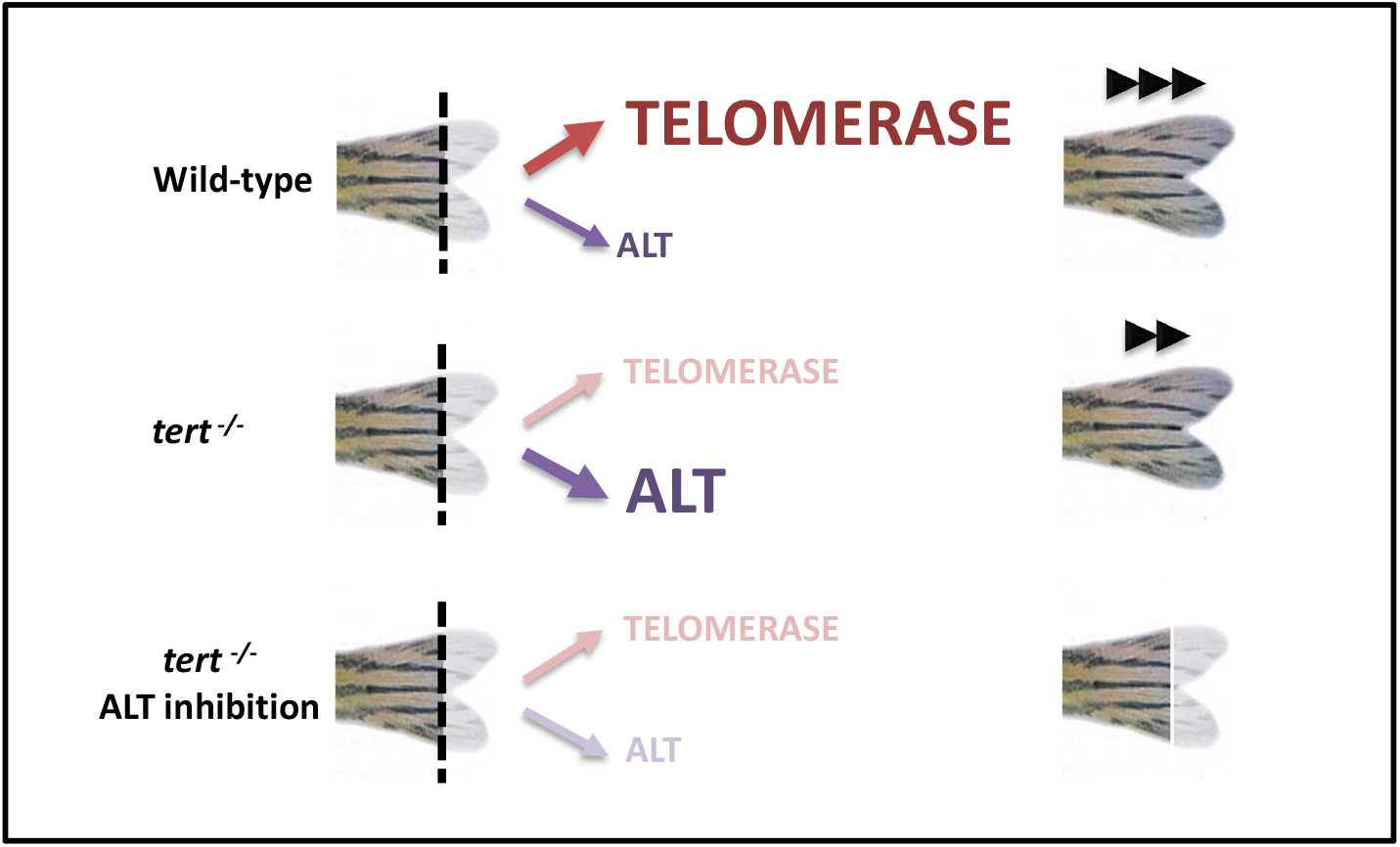
Schematic representation of telomeric maintenance in zebrafish caudal fin. Telomerase is the main telomeric maintenance mechanism that allows the regeneration of the tail fin after damage. However, in telomerase deficient fish, regeneration occurs thanks to ALT, although more slowly. Therefore, if both mechanisms are inhibited, no regeneration of zebrafish caudal fin is possible.

Therefore, ALT mechanism allows the regeneration of the tail fin in telomerase deficient zebrafish. However, not all zebrafish tissues can regenerate in the absence of telomerase, for example telomerase is essential for zebrafish heart regeneration (Aix et al, 2018; Bednarek et al, 2015), where that absence of telomerase impairs proliferation leading to the accumulation of DNA damage, and *tert* ^*-/-*^ zebrafish hearts acquire a senescent phenotype [Bednarek et al., 2015]. This suggests that telomere maintenance mechanisms may be specific of tissues and organs. In adition the coexistence of both telomere meintenance mechanisms is also specific, since a zebrafish model of cancer that uses ALT for telomere maintenance, reverted all ALT features when *tert* was over-expressed (Idilli et al, 2020a), as has been described before in human (Perrem et al, 1999). Further studies are needed to elucidate a possible evolutionary explanation to these differences. One hypothesis would be that the fins are external organ, crucial for the survival of the species and highly exposed to external damage and predators so the existence of both mechanisms ensure telomere maintenance in cells implicated in the regeneration after injury of an organ essential for fish survival.

In the present work, we confirmed the impressive capability of the zebrafish to heal and regenerate fin tissues after an injury, only limited by aging, even in the absence of telomerase, and the involvement of ALT mechanisms in the telomere length maintenance during the regeneration process of the zebrafish caudal fin. To our knowledge, this is the first tissue of an animal model demonstrating the coexistence of telomerase and ALT mechanism, only found in some cancer populations and engineered cellular models (De Vitis et al, 2018). Zebrafish has proven to be an exceptionally useful model for further studies on the role of telomeres and telomerase in regeneration.

On the other hand, ALT is only active in about 10% of tumors, but these are tumors with a poor prognosis. ALT mechanism is a potential target for therapy (Zhang & Zou, 2020). A zebrafish model of cancer that uses ALT for telomere maintenance was recently described (Idilli et al, 2020a; Idilli et al, 2020b). However, an ALT model of regeneration has never been described and would be a powerfull tool to test drugs. Zebrafish has proven to be an undisputably useful vertebrate model system, with great advantages over other models, such as the ease, cost and speed of screening compounds (Murphey & Zon, 2006; Yoganantharjah & Gibert, 2017). In this work, the validity of zebrafish larvae as a model for testing drugs that inhibits ALT and/ or telomerase, which can be used in large-scale screening, has been demonstrated.

## ACKNOWLEDGEMENTS

We warmly acknowledge Cynthia Cabello, María C. López-Maya, Inmaculada Fuentes and Pedro. J. Martínez for their excellent technical assistance. This work was supported by the Spanish Ministry of Science and Innovation (grants PI19/00188 to MLC), all co-funded with Fondos Europeos de Desarrollo Regional), Fundación Ramón Areces (grant to MLC), University of Murcia (predoctoral contract to EMB) and the University Catolic of Murcia (predoctoral contract to DHS). The funders had no role in study design, data collection and analysis, decision to publish, or preparation of the manuscript.

## MATERIALS AND METHODS

### Ethics Statement

The experiments performed comply with the Guidelines of the European Union Council (Directive 2010/63/EU) and the Spanish RD 53/2013 Experiments and procedures were performed as approved by the Bioethical Committee of the University Hospital “Virgen de la Arrixaca” (210621/4/ISCIII).

### Maintenance of zebrafish

Zebrafish (*Danio rerio* H., Cypriniformes, Cyprinidae) were obtained from the Zebrafish International Resource Center (ZIRC) and mated, staged, raised and processed using standard procedures. Details of husbandry and environmental conditions are available on protocols.io (DOI:dx.doi.org/10.17504/protocols.io.mrjc54n) (Widrick et al., 2018). *tert* mutant line (allele hu3430) was obtained from the Sanger Institute.

### Caudal fin regeneration assay on adult zebrafish

For the caudal fin regeneration assay, zebrafish from *tert* ^*+/+*^, *tert* ^*+/-*^ and *tert* ^*-/-*^ genotypes were anesthetized with 0.05% benzocaine (Sigma). Fin tissue was removed approximately 2mm from the base of the caudal peduncle using a razor blade. For the single amputation assay or the successive amputations assay, images of zebrafish fins were taken before and after the amputation, as well as from different days post-amputation (dpa) following the respective experimental design. The fin tissue was preserved and processed accordingly in each experiment. When the fish recovered, they were returned to recirculating water heated to 32°C for the duration of the experiment. Each fish was tracked and imaged individually to calculate regeneration progress over time. Percent fin regeneration was determined based on the area of regrowth divided by the original fin area.

### Cell isolation

Regenerated portion of caudal fin tissue obtained from zebrafish at different clips were incubated in Trypsin (0.5 mg/ml)/EDTA (0.1 mg/ml) (Gibco) in PBS for 1 minute (min), centrifuged (600 g, 5 min) and then incubated in Colagenase (0.5 mg/ml) (Gibco) in RPMI medium (Lonza) supplemented with CaCl_2_ 2H_2_O (0.7 mg/ml) for 30 min. The cell suspensions were obtained by pipetting, smashing and filtering the digested tissues through a 100 µm mesh and a 70 µm mesh (Becton Dickinson) successively, finally washed and resuspended in PBS.

### Flow-FISH

Cells from each fin sample obtained as described before were washed in 2 ml PBS supplemented with 0.1% bovine serum albumin (BSA) (NEB). Each sample was divided into two replicate tubes: one pellet was resuspended in 500 ml hybridization buffer and another in hybridization buffer without FITC-labeled telomeric PNA probe (PNABio) as negative control. Samples were then denatured for 10 min at 80°C under continuous shaking and hybridized for 2 h in the dark at room temperature. After that, the cells were washed twice in a washing solution (70% deionized Formamide (Sigma), 10 mM Tris (Sigma), 0.1% BSA (NEB) and 0.1% Tween-20 (Sigma) in dH_2_O, pH 7.2). The cells were then centrifuged, resuspended in 500 ml of propidium iodide solution, incubated for 2 hours at room temperature, stored at 4°C and analyzed by flow cytometry within the following 48 h.

### Q-FISH

Cell suspensions from regenerative tissue were obtained as described before and fixed in methanol:acetic acid (3:1). Interphases were spread on glass slides and dried overnight, and FISH was performed as described in Canela et al. (Canela et al, 2007). Cy3 and DAPI images were captured at 100x magnification using a confocal microscope (LSM 510 META from ZEISS, Jena, Germany) and Zeiss Efficient Navigation (ZEN) interface software. Telomere fluorescence signals were quantified using ImageJ software.

### C-Circles assay

The assay for the detection of telomeric DNA circles was performed as described (Lau et al, 2013) with modifications. Zebrafish caudal fin tissues collected from *tert* ^*+/+*^ and *tert* ^*-/-*^ genotypes at 0 hours post-amputation (hpa), and from the corresponding regenerate portion at 24 and 48 hpa were preserved at -80°C. Genomic DNA (gDNA) was extracted from the fin samples using the “ChargeSwitch™ gDNA Micro Tissue Kit (Invitrogen). A rolling circle amplification reaction (RCA) of partially double-stranded C-circles was performed with 0.2μg/µl bovine serum albumin (NEB), 0.1% Tween-20 (Sigma), 4μM dithiotreitol (DTT) (Invitrogen), 1 μM dNTPs (Invitrogen), 3.75U ɸ29 polymerase (NEB), 1X ɸ29 polymerase buffer (NEB) and 64 ng of gDNA. The mix was incubated at 30°C for 8 h and then at 65°C for 20 min. Finally, the RCA reaction was diluted to 40 μl with water. For each sample, the assay was done with and without ɸ29 polymerase, capable of amplifying circular DNA in a specific way.

After that, the telomeric secuences were detected through real-time PCR using C-circles amplification reaction as template. Real-time PCR was performed with a StepOnePlus instrument (Applied Biosystems) using SYBR® Premix Ex Taq™ (Perfect Real Time) (Takara). The primers (Telom F-Telom R) used are in Table 1 (Fig. S3). Finally, a second real-time PCR was performed with the same RCA to detect ribosomal protein s11 (*rps11*) as a single-copy gene used to normalize. The primers used are in Table 1 (Fig. S3). Both qPCRs were performed using the RCA with and without ɸ29 polymerase amplification as template. The C-circles amount value was obtained as follows: each telomeric qPCR was normalized to the corresponding *rps11* qPCR. Then, the normalized value of the samples without ɸ29 polymerase was subtracted from that with ɸ29 polymerase, and this final value was named as NORMTEL. The C-circles amount value is 2^NORMTEL^. As reference assay, the same protocol was performed using gDNA obtained from ALT-positive human osteosarcoma tumor cell line (Saos-2) and ALT-negative human cervical tumor cell line (HeLa).

### Gene expression analysis

Total RNA was extracted from the homogenized regenerative tissue in TRIzol Reagent (Ambion) and using the PureLink RNA Mini kit (Ambion), following the manufacturer’s instructions. RNA was treated with DNase I, amplification grade (1 U/mg RNA; Thermo Fisher Scientific). cDNA was generated by the SuperScript™ IV VILO™ Master Mix (Invitrogen), following the manufacturer’s instructions. Real-time PCR was performed with a StepOnePlus instrument (Applied Biosystems) using SYBR® Premix Ex Taq™ (Perfect Real Time) (Takara). Reaction mixtures were incubated for 30 seconds (sec) at 95°C, followed by 40 cycles of 5 sec at 95°C, 20 sec at 60°C, and finally a melting curve protocol. Ribosomal protein s11 (*rps11*) content in each sample was used for normalization of zebrafish mRNA expression, using the comparative *Ct* method (2^-ΔCt^). The primers used are shown in Table 1 (Fig. S3).

### Morpholino injection and electroporation

Adult *tert* ^*+/+*^ and *tert* ^*-/-*^ zebrafish were anesthetized in 0.05% benzocaine prior to the amputation of the distal portion of their caudal fins, proximal to the first lepidotrichial branching point. Following the surgery, the fish were returned to recirculating water heated to 32°C. Vivo morpholino oligonucleotides (MO) (Gene Tools) containing a 3’ fluorescein tag were injected at 1.5 mM into the regenerating tissue on the dorsal part of each zebrafish tail fin at 48 hpa. The other uninjected half (ventral area) was considered as an internal control in order to monitor the normal growth. Immediately after injections, the dorsal half was electropored using a NEPAGENE electroporator, five consecutive 50 msec pulses, at 15 V with a 30 sec pause between pulses, were used. A 3-mm-diameter platinum plate electrode (CUY 650-P3 Tweezers, Protech International) localized the pulses to approximately the dorsal one-half of the fin. The MO sequences are shown in Table 2 (Fig. S4).

After the procedure and 48 hours post-injection (hpi), fins were photographed capturing fluorescent and brightfield signals simultaneously using a Leica M205 FA fluorescence stereo microscope equipped with a DFC365FX camera (Leica). In order to calculate the percentage area of growth between the injected and non-injected part, the values were inserted in the following formula: (Dorsal 48 hpi - Dorsal 0 hpi)/(Ventral 48 hpi -Ventral 0 hpi)*100, where Dorsal is the regenerative area of the MO-treated tissue and Ventral is the regenerative area of the corresponding uninjected half.

### Amputation of zebrafish larval fin primordia and chemical exposure by immersion

48 hours post-fertilization (hpf) zebrafish larvae obtained from *tert* ^*+/-*^ breeding crosses were anesthetized in 0.008% tricaine (Sigma) and placed on an agar plate to amputate the caudal fin primordia with a surgical razor blade just posterior to the notochord. The larvae were individualized in each well of 96-well plates containing ATR Inhibitor IV (CAS 1232410-49-10 Calbiochem) at 10 µM or DMSO in egg water as negative control. Larval fin was imaged before amputation, after amputation and 24 hpa to measure and calculate the regeneration area in both conditions.

### Western blot

48 hpf zebrafish larvae were treated for 24 hours with 10 µM and 50 µM ATR Inhibitor IV (Calbiochem) or DMSO as negative control. The protein extracts were subjected to polyacrylamide gel electrophoresis, wet transferred to a nylon membrane (GE Healthcare) and analyzed by Western blot. The membrane was incubated for 1 h with TTBS (Tween-Tris-buffered saline) containing 5% (w/v) skimmed dried milk powder and immunoblotted using Phospho-(Ser/Thr) ATM/ATR Substrate Antibody (#2851 Cell Signaling) (dilution 1:1000) 16 h at 4°C. The blot was then washed with TTBS and incubated for 1 h at room temperature with the secondary HRP-conjugated antibody (dilution 1:1000) in 5% (w/v) skimmed milk in TTBS. After repeated washes, the signal was detected with the enhanced chemiluminescence reagent and ChemiDoc XRS (Biorad).

---

**Figure S1.**
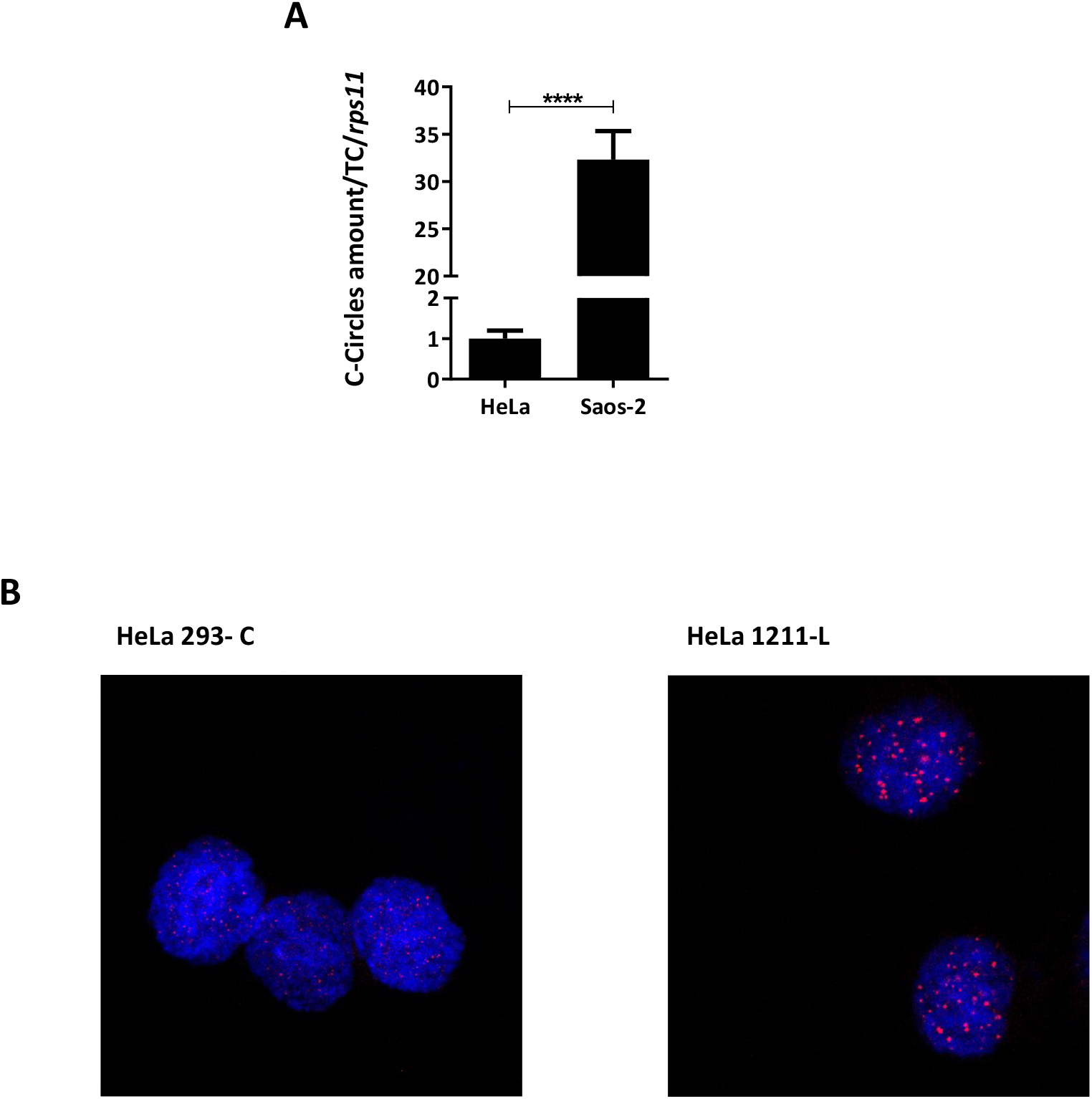
C-circles abundance and telomere QFISH in control cell lines. **A**, C-circles abundance in ALT-negative human cervical tumor cell line (HeLa) and ALT-positive human osteosarcoma tumor cell line (Saos-2). **B**. QFISH in HeLa cells. HeLa 293 has a telomere length of 2,7 kb and HeLa 1211 has a telomere length of 23 kb.

**Figure S2.**
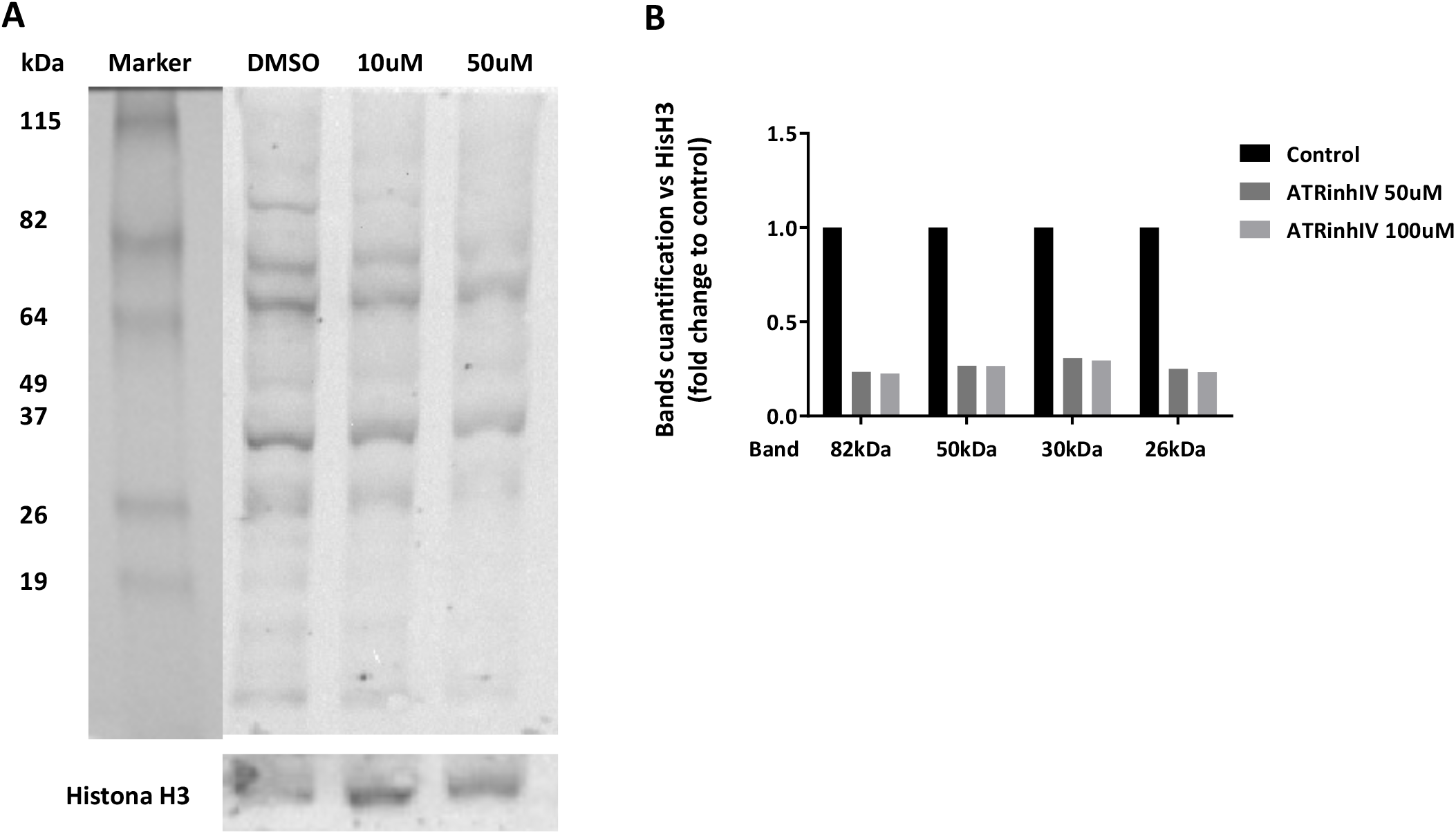
Determination of the effect of several concentrations of ATR Inhibitor IV on zebrafish larvae. **A**, Western blot against ATR/ATM substrates in 48 hpf zebrafish larvae treated for 24 hours with ATR Inhibitor IV at 10 µM and 50 µM. **B**, Quantification of various western blot bands against ATR/ATM substrates in relation to the histone H3 signal.

**Figure S3.**
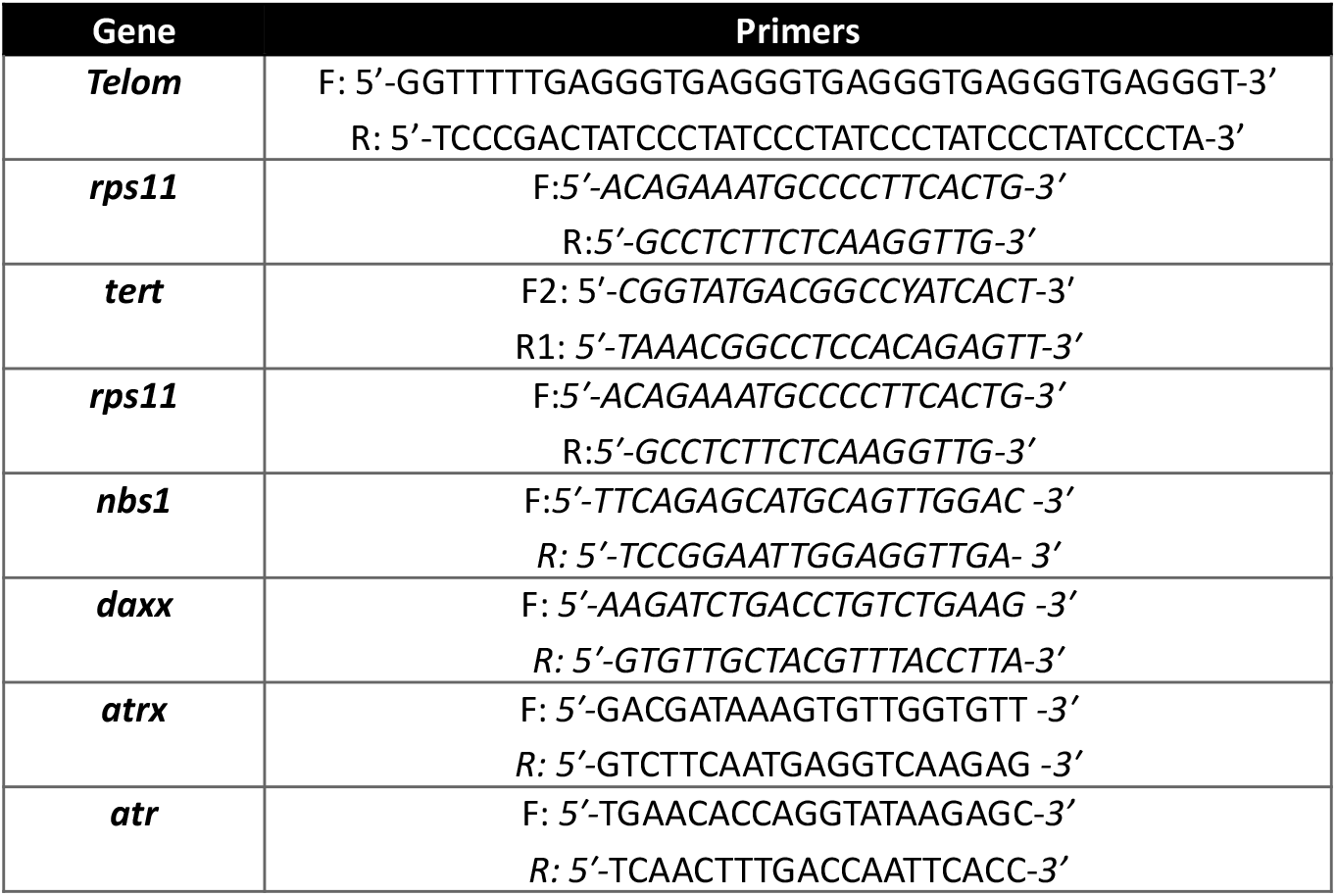
Primers used in this work.

**Figure S4.**
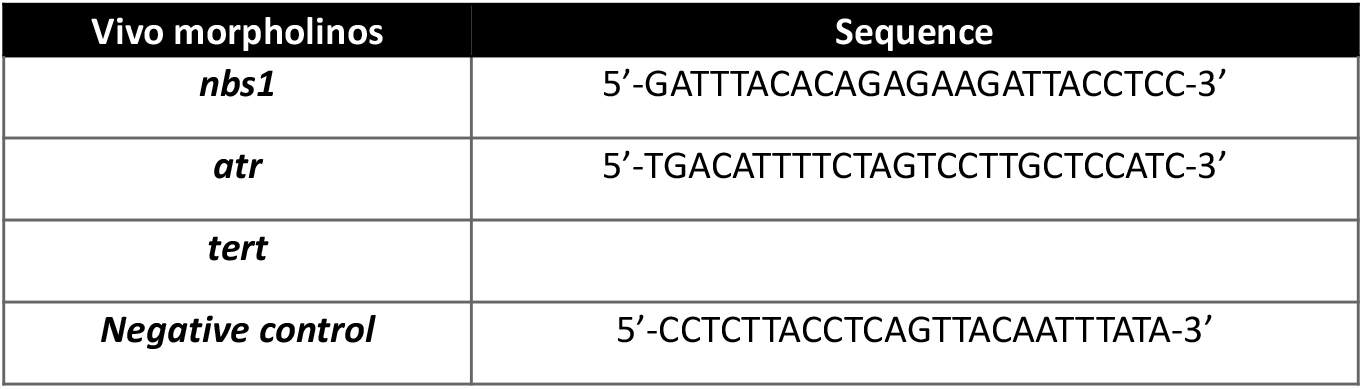
Vivo morpholinos used in this work.

## BIBLIOGRAPHY

Aix E, Gallinat A, Flores I (2018) Telomeres and telomerase in heart regeneration. Differentiation 100: 26–30

Akimenko MA, Mari-Beffa M, Becerra J, Geraudie J (2003) Old questions, new tools, and some answers to the mystery of fin regeneration. Developmental dynamics : an official publication of the American Association of Anatomists 226: 190–201

Amorim JP, Santos G, Vinagre J, Soares P (2016) The Role of ATRX in the Alternative Lengthening of Telomeres (ALT) Phenotype. Genes 7

Anchelin M, Alcaraz-Perez F, Martinez CM, Bernabe-Garcia M, Mulero V, Cayuela ML (2013) Premature aging in telomerase-deficient zebrafish. Disease models & mechanisms 6: 1101–1112

Anchelin M, Murcia L, Alcaraz-Perez F, Garcia-Navarro EM, Cayuela ML (2011) Behaviour of telomere and telomerase during aging and regeneration in zebrafish. PloS one 6: e16955

Azevedo AS, Grotek B, Jacinto A, Weidinger G, Saude L (2011) The regenerative capacity of the zebrafish caudal fin is not affected by repeated amputations. PloS one 6: e22820

Becker T, Wullimann MF, Becker CG, Bernhardt RR, Schachner M (1997) Axonal regrowth after spinal cord transection in adult zebrafish. The Journal of comparative neurology 377: 577–595

Bednarek D, Gonzalez-Rosa JM, Guzman-Martinez G, Gutierrez-Gutierrez O, Aguado T, Sanchez-Ferrer C, Marques IJ, Galardi-Castilla M, de Diego I, Gomez MJ, Cortes A, Zapata A, Jimenez-Borreguero LJ, Mercader N, Flores I (2015) Telomerase Is Essential for Zebrafish Heart Regeneration. Cell Rep 12: 1691–1703

Blasco MA, Lee HW, Hande MP, Samper E, Lansdorp PM, DePinho RA, Greider CW (1997) Telomere shortening and tumor formation by mouse cells lacking telomerase RNA. Cell 91: 25–34

Brault ME, Autexier C (2011) Telomeric recombination induced by dysfunctional telomeres. Mol Biol Cell 22: 179–188

Canela A, Vera E, Klatt P, Blasco MA (2007) High-throughput telomere length quantification by FISH and its application to human population studies. Proceedings of the National Academy of Sciences of the United States of America 104: 5300–5305

Cesare AJ, Griffith JD (2004) Telomeric DNA in ALT cells is characterized by free telomeric circles and heterogeneous t-loops. Molecular and cellular biology 24: 9948–9957

Collis SJ, Ciccia A, Deans AJ, Horejsi Z, Martin JS, Maslen SL, Skehel JM, Elledge SJ, West SC, Boulton SJ (2008) FANCM and FAAP24 function in ATR-mediated checkpoint signaling independently of the Fanconi anemia core complex. Mol Cell 32: 313–324

Compton SA, Choi JH, Cesare AJ, Ozgur S, Griffith JD (2007) Xrcc3 and Nbs1 are required for the production of extrachromosomal telomeric circles in human alternative lengthening of telomere cells. Cancer research 67: 1513–1519

Conomos D, Pickett HA, Reddel RR (2013) Alternative lengthening of telomeres: remodeling the telomere architecture. Frontiers in oncology 3: 27

De Vitis M, Berardinelli F, Sgura A (2018) Telomere Length Maintenance in Cancer: At the Crossroad between Telomerase and Alternative Lengthening of Telomeres (ALT). Int J Mol Sci 19

Draskovic I, Londono Vallejo A (2013) Telomere recombination and alternative telomere lengthening mechanisms. Front Biosci (Landmark Ed) 18: 1–20

Elmore LW, Norris MW, Sircar S, Bright AT, McChesney PA, Winn RN, Holt SE (2008) Upregulation of telomerase function during tissue regeneration. Exp Biol Med (Maywood) 233: 958–967

Fajkus J, Sykorova E, Leitch AR (2005) Techniques in plant telomere biology. Biotechniques 38: 233–243

Flynn RL, Cox KE, Jeitany M, Wakimoto H, Bryll AR, Ganem NJ, Bersani F, Pineda JR, Suva ML, Benes CH, Haber DA, Boussin FD, Zou L (2015) Alternative lengthening of telomeres renders cancer cells hypersensitive to ATR inhibitors. Science 347: 273–277

Gemberling M, Bailey TJ, Hyde DR, Poss KD (2013) The zebrafish as a model for complex tissue regeneration. Trends Genet 29: 611–620

Goldsmith JR, Jobin C (2012) Think small: zebrafish as a model system of human pathology. J Biomed Biotechnol 2012: 817341

Henriques CM, Carneiro MC, Tenente IM, Jacinto A, Ferreira MG (2013) Telomerase is required for zebrafish lifespan. PLoS Genet 9: e1003214

Henson JD, Cao Y, Huschtscha LI, Chang AC, Au AY, Pickett HA, Reddel RR (2009) DNA C-circles are specific and quantifiable markers of alternative-lengthening-of-telomeres activity. Nature biotechnology 27: 1181–1185

Hu Y, Shi G, Zhang L, Li F, Jiang Y, Jiang S, Ma W, Zhao Y, Songyang Z, Huang J (2016) Switch telomerase to ALT mechanism by inducing telomeric DNA damages and dysfunction of ATRX and DAXX. Scientific reports 6: 32280

Idilli AI, Cusanelli E, Pagani F, Berardinelli F, Bernabe M, Cayuela ML, Poliani PL, Mione MC (2020a) Expression of tert Prevents ALT in Zebrafish Brain Tumors. Front Cell Dev Biol 8: 65

Idilli AI, Pazzi C, Dal Pozzolo F, Roccuzzo M, Mione MC (2020b) Rad21 Haploinsufficiency Prevents ALT-Associated Phenotypes in Zebrafish Brain Tumors. Genes 11

Lau LM, Dagg RA, Henson JD, Au AY, Royds JA, Reddel RR (2013) Detection of alternative lengthening of telomeres by telomere quantitative PCR. Nucleic acids research 41: e34

Lovejoy CA, Li W, Reisenweber S, Thongthip S, Bruno J, de Lange T, De S, Petrini JH, Sung PA, Jasin M, Rosenbluh J, Zwang Y, Weir BA, Hatton C, Ivanova E, Macconaill L, Hanna M, Hahn WC, Lue NF, Reddel RR, Jiao Y, Kinzler K, Vogelstein B, Papadopoulos N, Meeker AK (2012) Loss of ATRX, genome instability, and an altered DNA damage response are hallmarks of the alternative lengthening of telomeres pathway. PLoS Genet 8: e1002772

Martinez AR, Kaul Z, Parvin JD, Groden J (2017) Differential requirements for DNA repair proteins in immortalized cell lines using alternative lengthening of telomere mechanisms. Genes, chromosomes & cancer 56: 617–631

McChesney PA, Elmore LW, Holt SE (2005) Vertebrate marine species as model systems for studying telomeres and telomerase. Zebrafish 1: 349–355

Morrison JI, Loof S, He P, Simon A (2006) Salamander limb regeneration involves the activation of a multipotent skeletal muscle satellite cell population. The Journal of cell biology 172: 433–440

Murnane JP, Sabatier L, Marder BA, Morgan WF (1994) Telomere dynamics in an immortal human cell line. EMBO J 13: 4953–4962

Murphey RD, Zon LI (2006) Small molecule screening in the zebrafish. Methods 39: 255–261

Napier CE, Huschtscha LI, Harvey A, Bower K, Noble JR, Hendrickson EA, Reddel RR (2015) ATRX represses alternative lengthening of telomeres. Oncotarget 6: 16543–16558

Perrem K, Bryan TM, Englezou A, Hackl T, Moy EL, Reddel RR (1999) Repression of an alternative mechanism for lengthening of telomeres in somatic cell hybrids. Oncogene 18: 3383–3390

Pickett HA, Reddel RR (2015) Molecular mechanisms of activity and derepression of alternative lengthening of telomeres. Nature structural & molecular biology 22: 875–880

Poss KD, Keating MT, Nechiporuk A (2003) Tales of regeneration in zebrafish. Developmental dynamics : an official publication of the American Association of Anatomists 226: 202–210

Poss KD, Nechiporuk A, Hillam AM, Johnson SL, Keating MT (2002) Mps1 defines a proximal blastemal proliferative compartment essential for zebrafish fin regeneration. Development 129: 5141–5149

Reimschuessel R (2001) A fish model of renal regeneration and development. ILAR journal 42: 285–291

Ren X, Tu C, Tang Z, Ma R, Li Z (2018) Alternative lengthening of telomeres phenotype and loss of ATRX expression in sarcomas. Oncology letters 15: 7489–7496

Rowlerson A, Radaelli G, Mascarello F, Veggetti A (1997) Regeneration of skeletal muscle in two teleost fish: Sparus aurata and Brachydanio rerio. Cell and tissue research 289: 311–322

Shao J, Chen D, Ye Q, Cui J, Li Y, Li L (2011) Tissue regeneration after injury in adult zebrafish: the regenerative potential of the caudal fin. Developmental dynamics : an official publication of the American Association of Anatomists 240: 1271–1277

Wehner D, Weidinger G (2015) Signaling networks organizing regenerative growth of the zebrafish fin. Trends Genet 31: 336–343

Yoganantharjah P, Gibert Y (2017) The Use of the Zebrafish Model to Aid in Drug Discovery and Target Validation. Current topics in medicinal chemistry 17: 2041–2055

Yost KE, Clatterbuck Soper SF, Walker RL, Pineda MA, Zhu YJ, Ester CD, Showman S, Roschke AV, Waterfall JJ, Meltzer PS (2019) Rapid and reversible suppression of ALT by DAXX in osteosarcoma cells. Scientific reports 9: 4544

Zhang JM, Zou L (2020) Alternative lengthening of telomeres: from molecular mechanisms to therapeutic outlooks. Cell Biosci 10: 30

Zhong ZH, Jiang WQ, Cesare AJ, Neumann AA, Wadhwa R, Reddel RR (2007) Disruption of telomere maintenance by depletion of the MRE11/RAD50/NBS1 complex in cells that use alternative lengthening of telomeres. The Journal of biological chemistry 282: 29314–29322

